# Escaping antibody responses to bacteriolytic enzymes Pal and Cpl-1 by epitope scanning and engineering

**DOI:** 10.1101/2022.10.06.511193

**Authors:** Marek Harhala, Katarzyna Gembara, Izabela Rybicka, Zuzanna Kaźmierczak, Paulina Miernikiewicz, Joanna Majewska, Wiktoria Budziar, Anna Nasulewicz-Goldeman, Daniel C. Nelson, Barbara Owczarek, Krystyna Dąbrowska

**Affiliations:** Hirszfeld Institute of Immunology and Experimental Therapy, Wroclaw, 53-114, Poland; Regional Specialist Hospital, Research and Development Centre, Wroclaw, 51-124, Poland; Institute for Bioscience and Biotechnology Research, University of Maryland, Rockville, 20850, Maryland, USA

**Keywords:** deimmunization, endolysin, Pal, Cpl-1, epitope engineering

## Abstract

Bacteriolytic enzymes are promising antibacterial agents, but they can cause a typical immune response *in vivo*. In this study, we used a targeted modification of two antibacterial endolysins, Pal and Cpl-1. We identified the key immunogenic amino-acids, and designed and tested new, bacteriolytic variants with altered immunogenicity. One new variant of Pal (257-259 MKS → TFG) demonstrated decreased immunogenicity while a similar mutant (257-259 MKS → TFK) demonstrated increased immunogenicity. A third variant (280-282 DKP → GGA) demonstrated significantly increased antibacterial activity and it was not cross-neutralized by antibodies induced by the wild-type enzyme. We propose this variant as a new engineered endolysin with increased antibacterial activity that is capable of escaping cross-neutralization by antibodies induced by wild-type Pal. We show that efficient antibacterial enzymes that avoid cross-neutralization by IgG can be developed by epitope scanning, *in silico* modelling, and substitutions of identified key amino acids with a high rate of success. Importantly, this universal approach can be applied to many proteins beyond endolysins and has the potential for design of numerous biological drugs.

## INTRODUCTION

Bacteriophage (or phage)-encoded cell wall hydrolases, termed endolysins, are currently being developed as novel therapeutic tools for treating infections by Gram-positive pathogens. Endolysins are normally active late in the phage infection cycle, rapidly cleaving bonds necessary for peptidoglycan stability in order to facilitate bacterial lysis and release of progeny phage. When applied directly to susceptible bacteria in a purified form in the absence of bacteriophage, endolysins act “from without” to hydrolyze the peptidoglycan, resulting in osmotic lysis of the organism (Nelson et al., 2012; Schmelcher & Loessner, 2021). Significantly, due to their rapid lytic actions, endolysins are not susceptible to efflux pumps, penicillin-binding proteins, alterations of metabolic pathways, or other mechanisms of resistance seen with traditional antibiotics, making them ideal alternative therapeutics to treat multi-drug resistant organisms (Gutiérrez et al., 2018; Oliveira et al., 2018). Additionally, endolysins are species-specific and, as such, represent a potential narrow-spectrum antimicrobial therapeutic that exploits the targeted killing capability of phage. Indeed, the US FDA granted “Breakthrough Therapy” designation status to endolysins in 2020 and several biotechnology/pharmaceutical companies have been conducting human clinical trials on endolysins, thus validating endolysin technologies as antimicrobial biologics, as reviewed (Murray et al., 2021).

Among the most notable endolysins in pre-clinical development are Pal, derived from *Streptococcus* phage Dp-1, and Cpl-1, derived from *Streptococcus* phage Cp-1. *Streptococcus pneumoniae*, the target of Cpl-1 and Pal, is the most common cause of bacteremia, pneumonia, meningitis, and otitis media in children - despite a successful vaccine campaign against pneumococcal disease over the past two decades, a Global Burden of Disease Study suggested that over 500,000 deaths still occur annually due to *S. pneumoniae* infection (Naghavi et al., 2015). Thus, pneumococcal endolysins represent a high potential for therapeutic application.

In spite of being promising antibacterial agents, endolysins are prokaryotic proteins that induce a normal immune response *in vivo* (Harhala et al., 2018; Jun et al., 2017; Schmelcher & Loessner, 2021). It has been demonstrated for lysostaphin, which is also a peptidoglycan-targeting antibacterial enzyme, that reduction of its immunogenicity translated into improved efficacy *in vivo* (Zhao et al., 2015). An engineered lysostaphin variant provided protection against repeated challenges of methicillin-resistant *Staphylococcus aureus* (MRSA), whereas the wild-type (WT) enzyme was efficacious only against the initial MRSA infection, but failed to clear subsequent bacterial challenges that were coincident with escalating anti-drug antibody titers. As demonstrated, endolysin-specific serum from immunized mice also reduced the rate of antibacterial activity of these enzymes (Harhala et al., 2018; Rashel et al., 2007). However, only general observations about the immunogenicity of endolysins has been made to date. Particularly, molecular features (i.e. epitopes defined by amino acid composition) that contribute to this effect remain unknown.

Although many investigators have developed improved endolysins with the aim of achieving better efficacy, these investigations only focus on improvements of endolysin stability or alterations of their bacterial host range or catalytic action. While these features are of great importance, they only relate to the basic activity that the enzyme exhibits *in vitro*. For *in vivo* applications, antibacterial drugs must overcome specific conditions and pressure from the living system, including the prominent effects of the immune system. In this study, we seek to understand how immune-reactive epitopes are distributed within endolysin proteins, and what modifications result in variants capable of escaping from the specific immune response induced by the WT enzymes. We apply true-epitope identification using the EndoScan platform technology developed here that allow us to identify epitopes that have elicited specific antibodies either in animals or in humans. Our approach applies, for the first time, a high-throughput method of epitope identification in endolysins. As such, it validates a novel, highly efficient method for improvement and adaptation of many other protein-based biological drugs and extends the existing methods of protein de-immunization by engineering identified antigenic determinants.

## RESULTS

### EndoScan: identification of antigenic epitopes in Pal and Cpl-1 endolysins

Identification of antigenic epitopes was done by epitope scanning (EndoScan), a new approach proposed herein for protein epitope identification, derived from the innovative VirScan technology developed by Xu et al. (Xu et al., 2015). The EndoScan approach is presented in Figure 1. Briefly, a phage display library of all epitopes in the endolysins was constructed with a nucleotide-printed library of oligopeptides representing overlapping sequences of the investigated endolysins. This library was incubated with (i) murine sera containing high levels of anti-Pal or anti-Cpl-1 IgG induced by specific challenge, or (ii) human sera representing the normal population of healthy volunteers.

**Figure 1.**
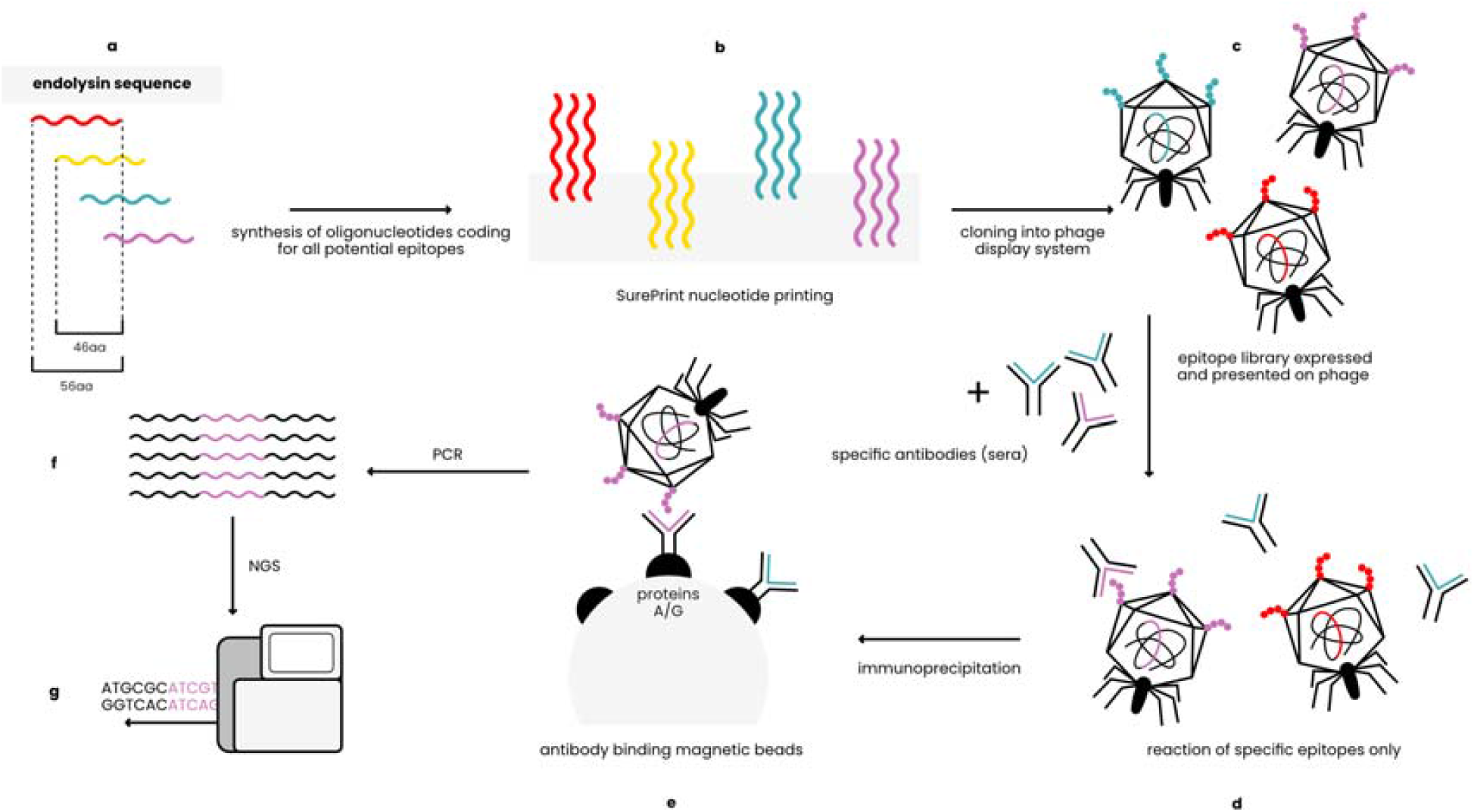
EndoScan technology – an overview (modified from Xu et al) (Xu et al., 2015). A, *In silico* design of peptides covering sequences of Pal and Cpl-1. B, Synthesis of oligonucleotides coding for the peptides. C, Constructing the phage display library of endolysin-derived peptides. D, Reaction of the library with endolysin-specific sera. E, Immunoprecipitation with magnetic beads binding Fc fragments of antibodies. F, Amplification by PCR reaction. G, NGS sequencing.

The investigated sera were immunoprecipitated with IgG-specific magnetic beads. Thus, a fraction of the phage display library that exposed endolysin epitopes reactive to IgG present in sera was isolated. Since the library contains whole phage particles, each phage that displays a specific epitope contains a sequence coding for this epitope in its genome. Thus, NGS sequencing of the relevant region in the library genomes was used to reveal which epitopes were effectively recognized by specific IgG (Figure 1).

Oligopeptides used in the library were 56 aa long and they overlapped each other by 46 aa, thus creating up to 6-fold coverage for any given residue, to avoid splitting epitopes or loosing interactions with surrounding peptide structures. Two types of these libraries were used to allow for complete analysis of immunogenic epitopes. The first library allowed for general screening and detection of immunoreactive regions within the endolysins; the same regions were identified in a mouse model and in a human population (Figure 2, Expanded View Table EV1).

**Figure 2.**
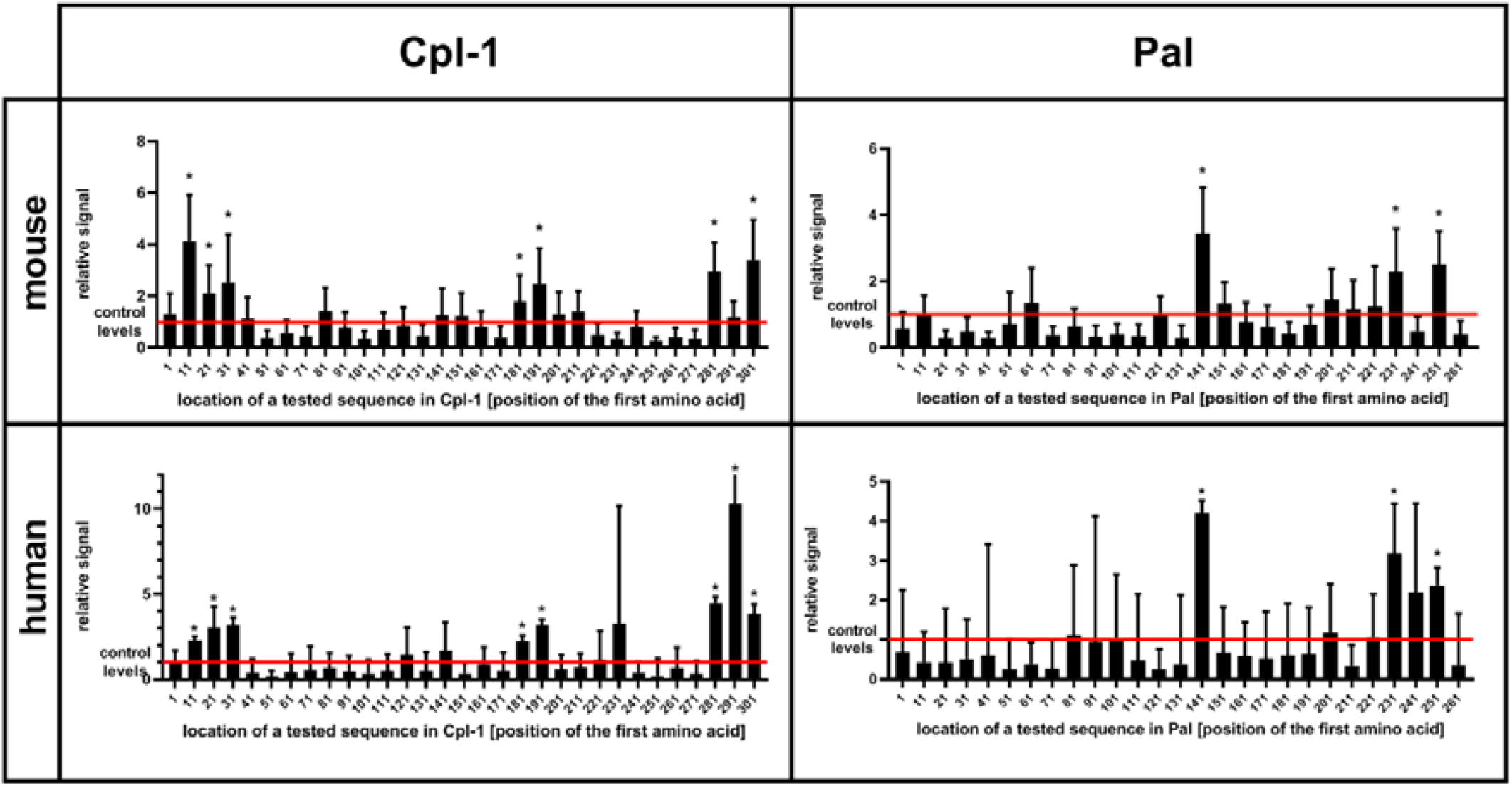
Immunoreactivity of short oligopeptides representing whole sequences of Cpl-1 and Pal endolysins in humans and mice. Oligopeptides used in the library were 56 aa long each and they overlapped by 46 aa (10 aa shift). Significantly, an increased relative signal (in comparison to control levels of input sample before immunoprecipitation) demonstrates interactions between IgG and the oligopeptide. X-axis specifies the position of the first amino acid in each oligopeptide as numbered in the WT protein. Oligopeptides were expressed as a phage display library, immunoprecipitated with sera from mice or humans representing a normal population of healthy volunteers. Data information: * adj. p value<0.05 between relative signal (ratio) of a measured oligopeptide after immunoprecipitation and before (control level in input sample, red line), Kruskal-Wallis test (two-sided), GraphPad Prism 9. Red line – levels for control group, a library before immunoprecipitation (input sample), data represent 7 (mouse) or 56 (human) biological replicates. Bars and whiskers represent mean value and standard deviation.

The second library was designed to define key amino acids forming the epitope sites. Specifically, when key amino acids are substituted, the variants fail to react with specific antibody (Expanded View Figure EV2). Thus, the second library contained a set of oligonucleotides coding for the same fragments of the endolysins, but bearing alanine substitutions in the sequence. Where the original sequence contained alanine, a glycine was used as the substitution. Further, to reduce the risk of false positive and false negative responses, variants with double substitutions and triple substitutions (targeted amino acid plus neighboring amino acids) were also included, resulting in 165 variants for each oligopeptide. The mouse sera was used for identification of amino acids that contributed to the Pal and Cpl-1-specific responses. The EndoScan technology as described above allowed for identification of 41 amino acids in Pal and 21 amino acids in Cpl-1 (Figure 3).

**Figure 3.**
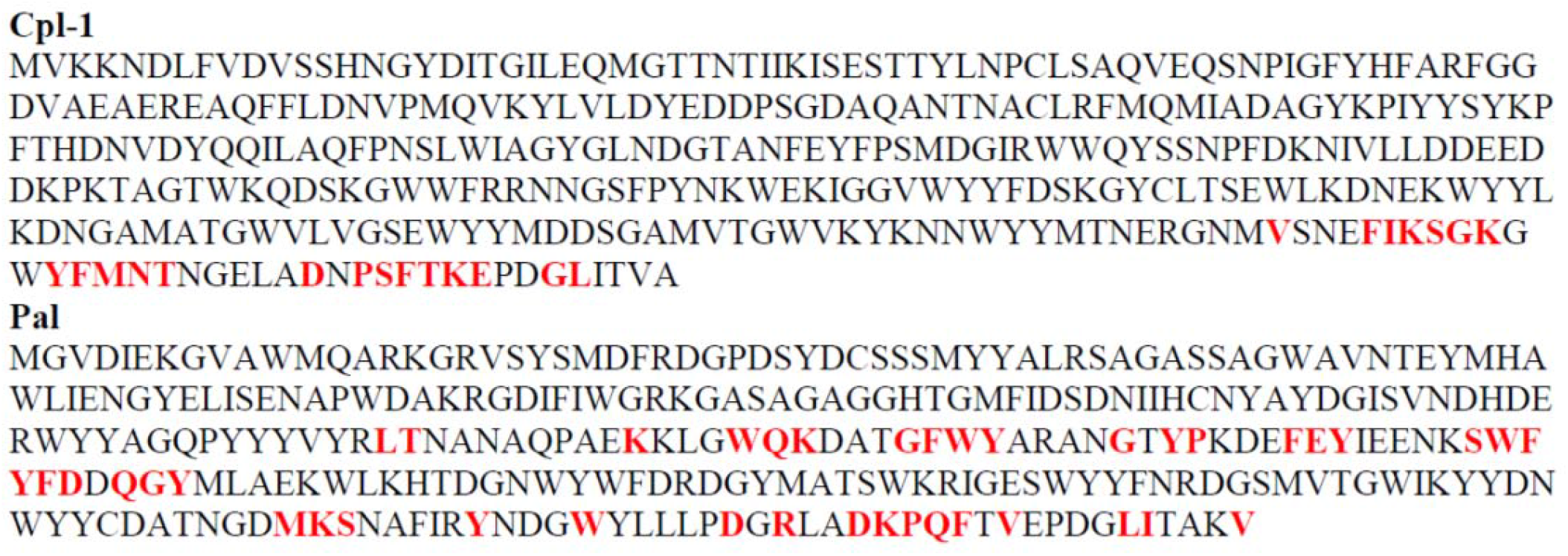
Key immunogenic amino acids (in red and bold) forming immunoreactive epitopes within Pal and Cpl-1 endolysins as defined by the EndoScan technology. Each amino acid in detected immunoreactive fragments was substituted with alanine (alanine with a glycine). If an oligopeptide after alanine substitution lost its previous immunoreactivity, such amino acid was designated as a key one for immunoreactivity of a protein.

### Pal and Cpl-1 variants with engineered immunoreactive epitopes

Engineering Pal and Cpl-1 to escape recognition by epitope-specific antibodies was conducted by amino acid substitutions with the following assumptions:

- each substitution was complemented with amino acids with different charge and chemical type of its substituents to disturb the epitope match with antibodies;
- the change in a folding energy of the protein (ΔΔG) calculated after substitution should be as low as possible for stabilization of the protein tertiary structure;
- smaller amino acids were preferred to larger amino acids to minimize steric tensions.

Nine possible variants of Pal were designed, but expression in the *E. coli* system yielded only three that were expressed as soluble proteins: Pal v1, Pal v3, and Pal v9. For Cpl-1, five possible variants were designed and they all yielded satisfactory expression of soluble enzymes (Table 1, full sequences are presented in Expanded View Figure EV3).

**Table 1.** Designed Pal and Cpl-1 variants with engineered immunoreactive epitopes. A list of modifications to WT endolysins tested experimentally.

| protein | variant designation | position of a modification | WT sequence | modified sequence |
| --- | --- | --- | --- | --- |
| Pal | v1 | 257-259 | MKS | TFK |
|  | v2 | 265-268 | YNDG | DKGL |
|  | v3 | 280-282 | DKP | GGA |
|  | v4 | 283-285 | QFT | GLA |
|  | v5 | 257-259<br>265-268<br>280-285 | MKS<br>YNDG<br>DKPQFT | AAA<br>AAAA<br>AAAAAA |
|  | v6 | 257-259<br>265-268<br>280-285 | MKS<br>YNDG<br>DKPQFT | TFK<br>DKGL<br>GGAGLA |
|  | V7 | 257-259<br>280-282 | MKS<br>DKP | TFK<br>GGA |
|  | v8 | 265-268 | YNDG | LAGL |
|  | v9 | 257-259 | MKS | TFG |
| Cpl-1 | v1 | 309-311 | SGK | DAW |
|  | v2 | 316 | M | R |
|  | v3 | 329-331 | TKE | LQP |
|  | v4 | 309-311<br>316-317<br>329-331 | SGK<br>MN<br>TKE | DAW<br>RF<br>LQP |
|  | v5 | 309-339 | SGK...TVA | deleted |

All expressed variants of Pal and Cpl-1 were tested for their antibacterial activity against pneumococci. This testing was completed over a range of enzyme concentrations and revealed that all selected variants had antibacterial activity, although the antibacterial activity in variants v1 and v9 of Pal and variants v2, v4, and v5 of Cpl-1 was significantly lower than in the WT enzymes (Figure 4). Variants Cpl-1 v1 and Cpl-1 v3 were active similarly to the WT enzyme, and Pal v3 was significantly more active against sensitive bacteria than the WT enzyme (Figure 4). Since all variants demonstrated at least partial antibacterial activity, they were all used in further immunological studies.

**Figure 4.**
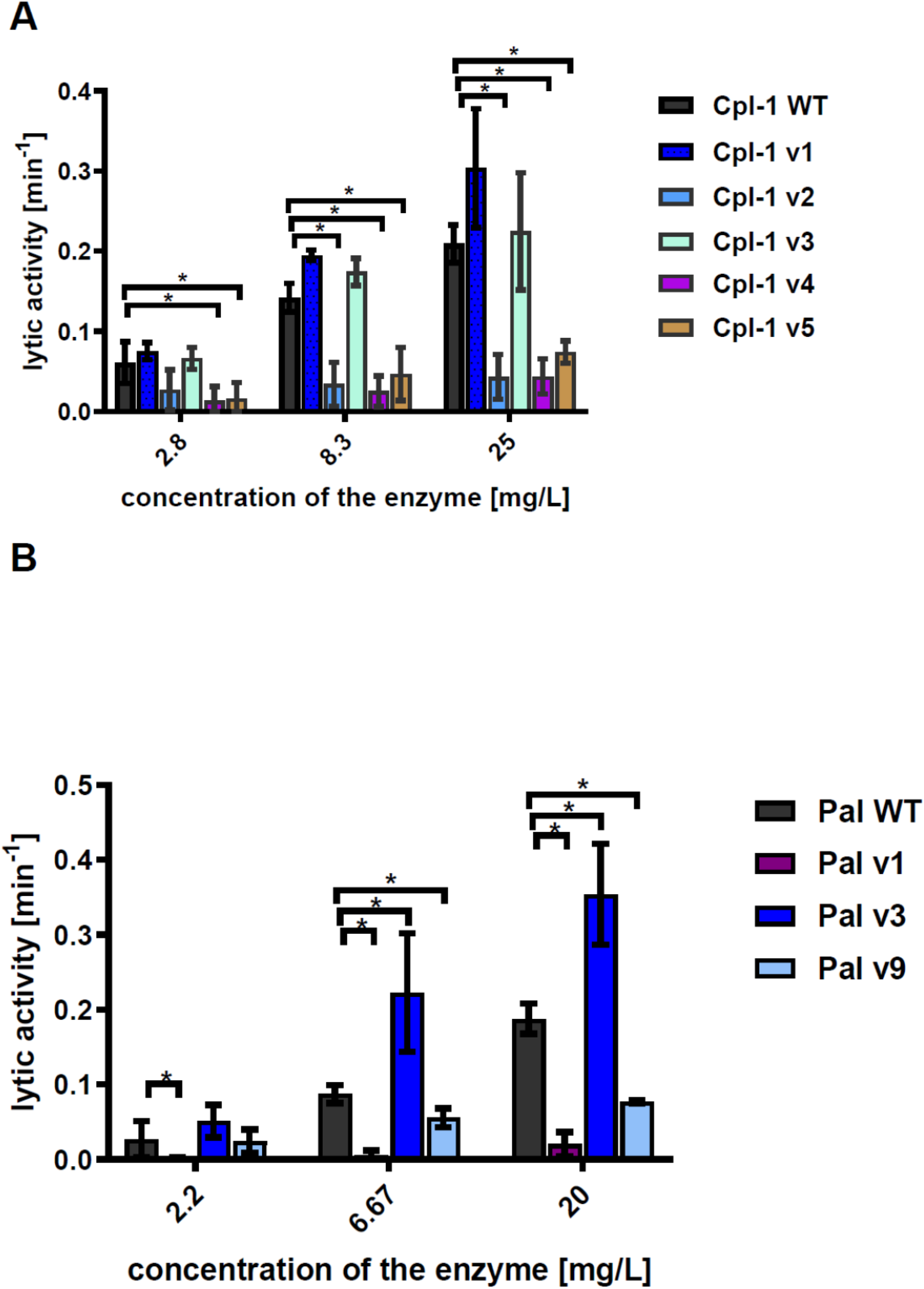
Antibacterial activity of variants in PBS. A, Antibacterial activity of endolysins Cpl-1 and its variants with engineered epitopes. B, Antibacterial activity of endolysins Pal and its variants with engineered epitopes. Data information: *Lytic activity* represents a part of the sample that was lysed in one minute at the peak activity. Bars and whiskers represent mean and SD of 3 experiments (two replicates each). * adj. *p*<0.05, unpaired, two-sided *t*-test with *p* value adjusted using the Holm-Sidak method, GraphPad Prism 9.

### Immune responses to Pal and Cpl-1 variants with engineered immunoreactive epitopes

Biologically active proteins can be prone to non-specific inactivation by the complex serum matrix. This might change the variants’ applicability *in vivo*, independent of their possible improved performance in the presence of antibodies targeting the WT endolysin. Endolysin variants were therefore tested for antibacterial activity *ex vivo* in naïve sera from unchallenged animals. In the WT enzymes, as well as most of the tested variants, sera were found to significantly decrease lytic activity (Figure 5). Three Cpl-1 variants, Cpl-1 v2, Cpl-1 v4 and Cpl-1 v5 demonstrated severely impaired overall activity, even though the Cpl-1 v4 did not demonstrate a significant difference in its activity with or without serum. Pal v1, also with intrinsic poor activity, demonstrated improved activity in the presence of serum, but it was still lower than that of the WT protein (Figure 5).

**Figure 5.**
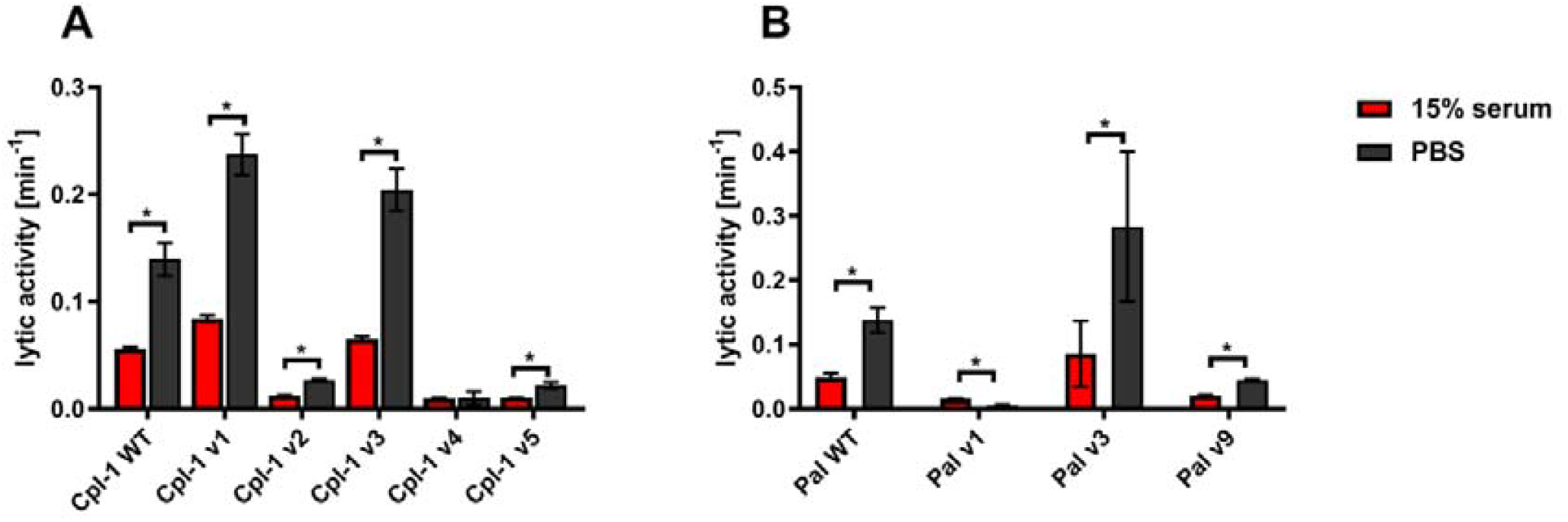
Effect of naïve (non-specific) murine serum on antibacterial activities of endolysins Cpl-1 and Pal. A, Comparison of activity in non-specific serum (15%) and PBS of Cpl-1 endolysin (10 μg/mL) and its variants. B, Comparison of activity in non-specific serum (15%) and PBS of Pal endolysin (10 μg/mL) and its variants. Data information: *Lytic activity* represents a part of the sample that was lysed in one minute at the peak activity. Bars and whiskers represent mean and SD of 3 experiments (two replicates each). * adj. *p*<0.05, unpaired, two-sided *t*-test with *p* value adjusted using the Holm-Sidak method, GraphPad Prism 9.

Pal and Cpl-1 variants were further investigated for their *in vivo* immunogenicity in an animal model. Immunogenicity herein is considered as an induction of serum levels of specific IgG. In the case of Pal v9, its immunogenicity was significantly decreased, but only on days 40 and 50 after challenge (*p*<0.001). In contrast, the immunogenicity of Pal v1 was significantly increased on days 40 and 50 after challenge (*p*<0.001) (Figure 6). Thus, the overall ability of the variants to induce specific antibodies in comparison to WT enzymes was decreased only in one case, and this variant demonstrated considerably lower antibacterial activity than the WT endolysin.

**Figure 6.**
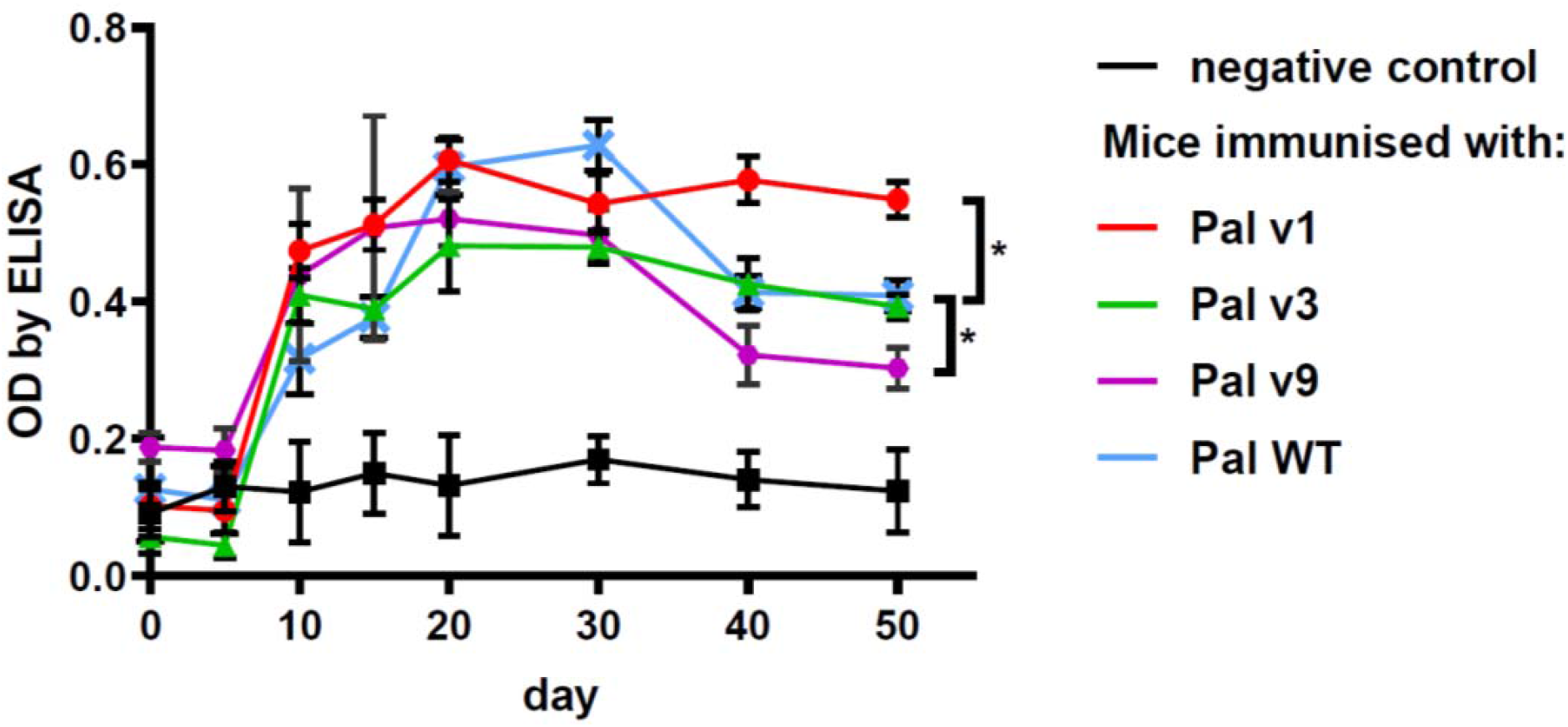
Serum levels of specific IgG induced by Pal and its variants with engineered epitopes identified by EndoScan. Data information: Points and whiskers represent mean and standard deviation of 6 biological replicates in a group, * - p<0.05 between indicated groups at a day 50 (biological replicates in each group, n=6), unpaired, two-sided *t*-test with *p* value adjusted using the Holm-Sidak method, GraphPad Prism 9.

Specific antibodies induced by an active protein can potentially limit success of this protein in situations of repeated use. Here, in most cases, we did not achieve overall deimmunization. Nonetheless, variants with the same overall immunogenicity but different immunogenic epitopes may escape cross-reactions with antibodies induced by the WT protein, thus being useful in repeat treatments. Therefore, we investigated by ELISA cross-reactions of Pal and Cpl-1 variants with IgG induced by the WT enzymes. In all variants, cross-reactivity was weaker than the reactivity of WT Pal or Cpl-1, with the decrease of the Pal variants’ cross-reactivity being more marked (Figure 7). Therefore, we next tested *ex vivo* how the Pal variants escaped cross-neutralization by specific serum induced by WT Pal. The lytic activity of Pal WT, and its variants v3 and v9, was compared in the reaction with Pal-specific serum used as a blocking agent. Pal v1 was not investigated due to its overall significantly weakened antibacterial activity. This testing revealed that Pal v3 demonstrated significantly stronger antibacterial activity in Pal-specific serum than the WT Pal enzyme (*p*<0.05). Specifically, two effects contributed to this advantage: Pal v3 was not neutralized by Pal-specific serum, and it demonstrated intrinsic higher activity than WT Pal (Figure 8). Thus, Pal v3 is a variant of the Pal endolysin that escapes cross-neutralization with specific antibodies induced by the WT Pal, and it has overall improved antibacterial efficacy.

**Figure 7.**
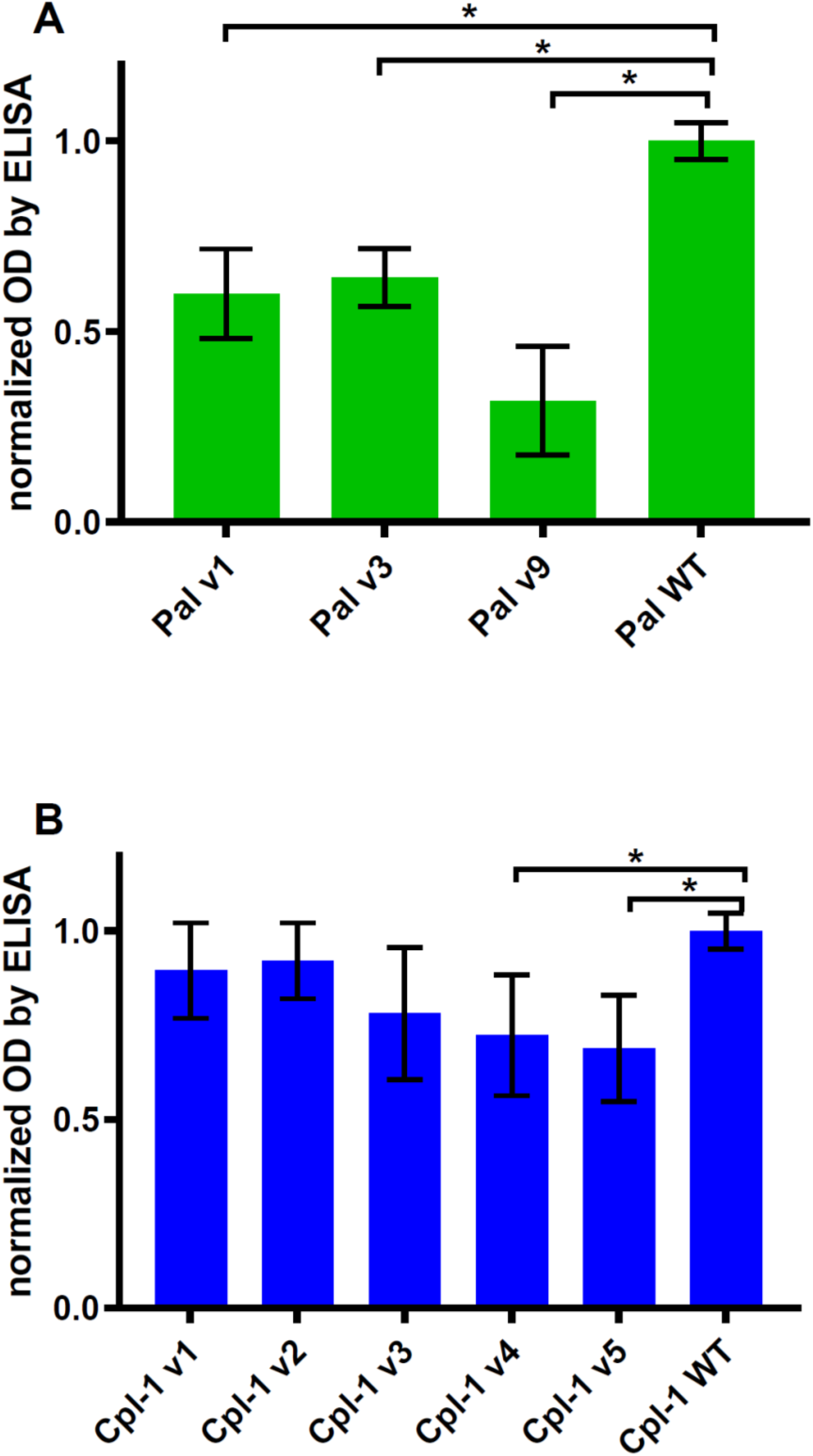
Cross-reactivity of serum with IgG specific to WT Pal and Cpl-1 to its variants. A, Cross-reactivity of WT Pal specific IgG. B, Cross-reactivity of WT Cpl-1 specific IgG. Data information: Bars and whiskers represent mean and standard deviation of 6 biological replicates per group. * adj. *p*<0.05, unpaired, one-sided *t*-test with *p* value adjusted using the Benjamini & Hochberg method (detailed information on statistics with *p* values is given in Expanded View Table EV2).

**Figure 8.**
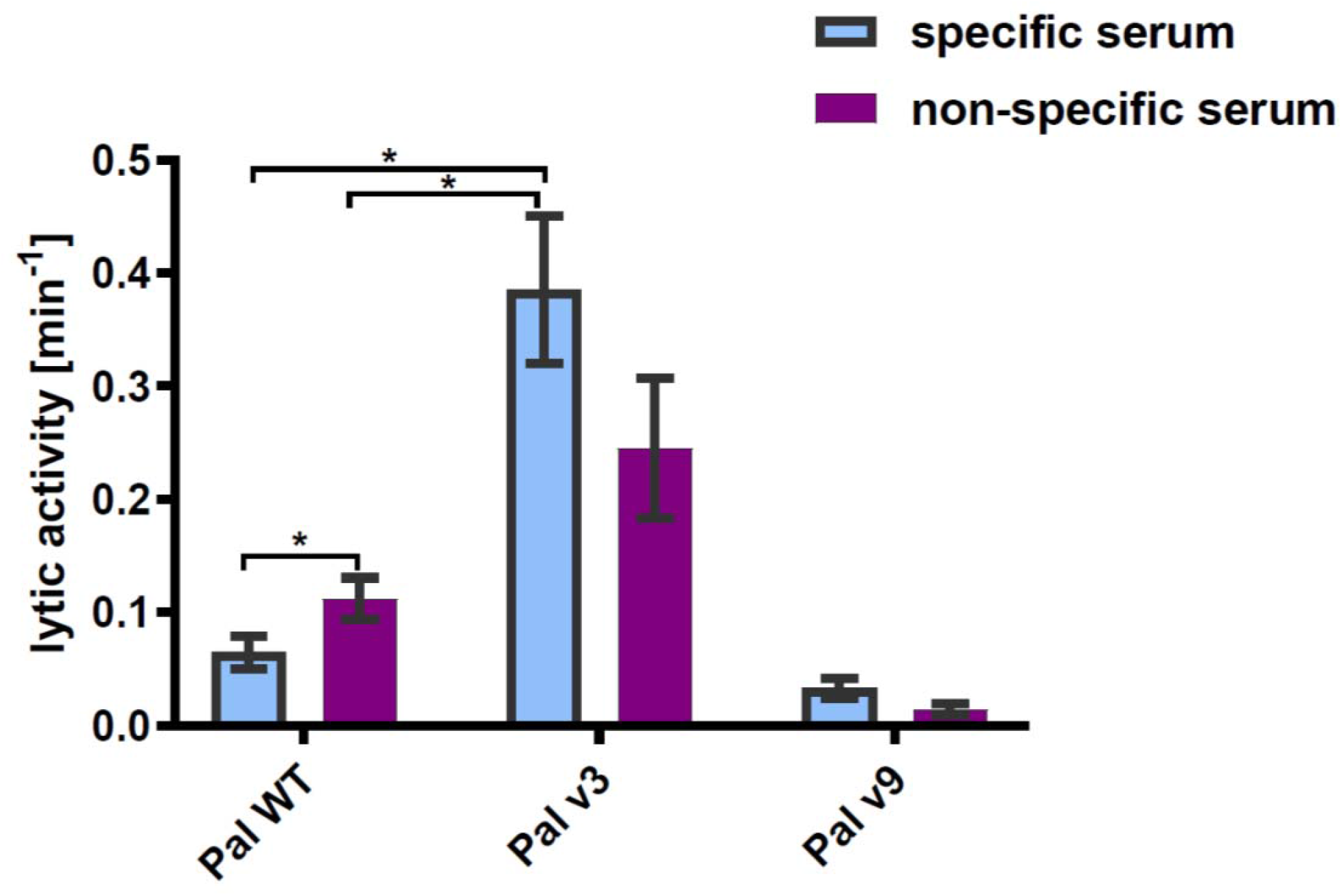
Lytic activity of WT Pal and its variants v3 and v9 in specific and non-specific serum. Data information: The tested concentration of endolysins and all variants was 20 μg/mL. Serum concentration was 15%. Bar and whiskers represent mean and SD of 3 measurements (two technical replicates each). * adj. *p*<0.05, unpaired *t*-test with *p* value adjusted using the Holm-Sidak method, GraphPad Prism 9.

**Figure 9:**
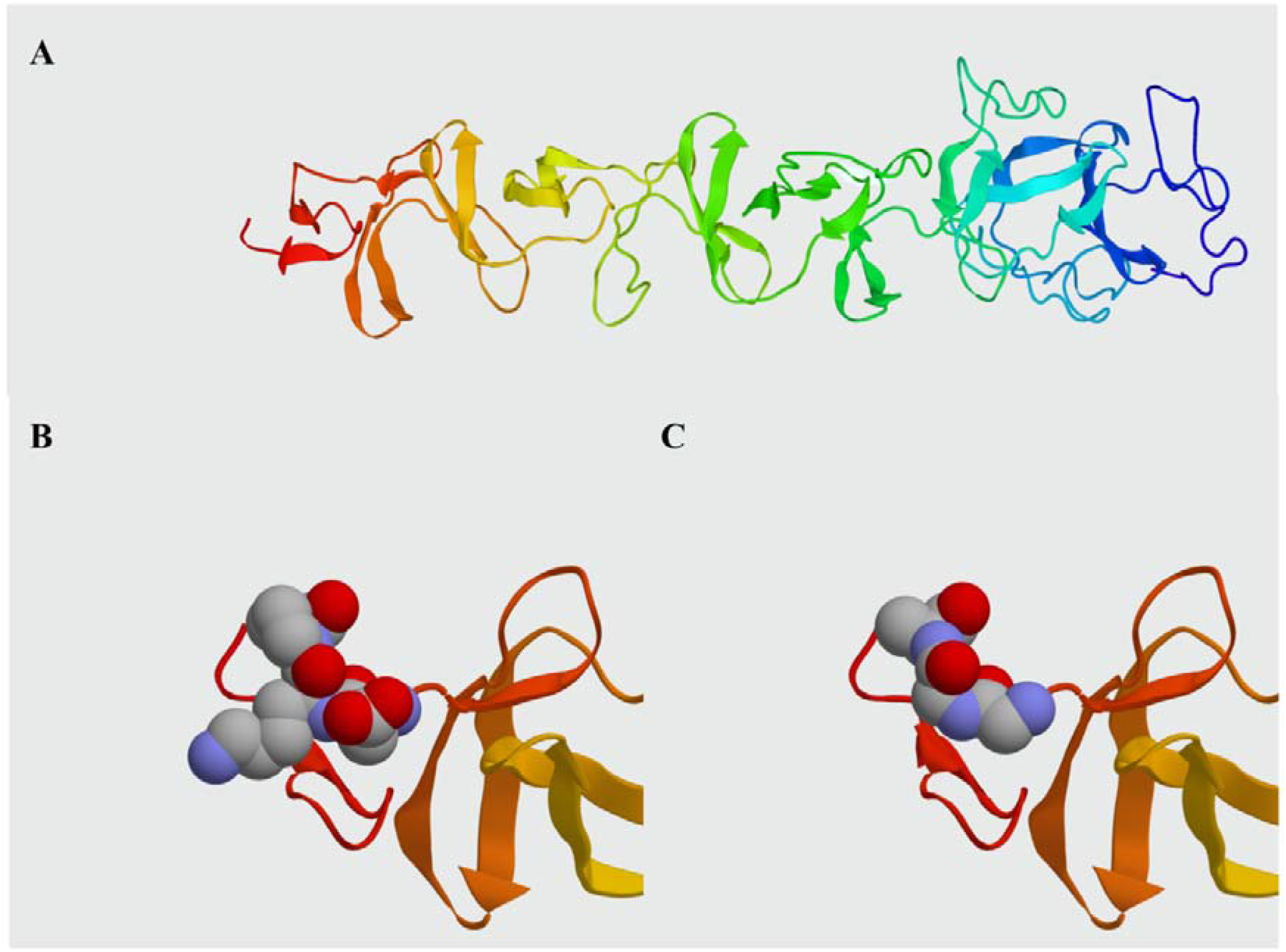
Comparison between Pal WT and Pal v3 (280-282: DKP→GGA) molecular structures. A, *In silico* model of Pal WT developed in I-TASSER. B, Presentation of epitope DKP in Pal WT. C, Presentation of epitope GGA replacing DKP in Pal v3. Data information: Atoms within the epitopes are represented by spheres: grey – carbon, red – oxygen, blue – nitrogen. Representation generated by Protean 3D software.

## DISCUSSION

In spite of being promising antibacterial agents, endolysins are prokaryotic proteins that have been demonstrated to induce a normal immune response *in vivo*, including specific antibodies induction (Harhala et al., 2018; Jun et al., 2017). Importantly, antibodies targeting a protein do not enter random interactions with the proteins surface. Rather, they bind only selected sites that constitute antigenic epitopes that mediate protein immunogenicity (Janeway, Charles A et al., 2001). In this study, we applied a modified approach of antigenic epitope scanning adapted from Xu et al. for identification of antigenic epitopes within two antibacterial enzymes, the endolysins Pal and Cpl-1 (Figure 2) (Xu et al., 2015). Scanning included both a mouse model and human sera. Animal sera provided a source of IgG specifically targeting the investigated endolysins, developed by challenge with WT Pal or Cpl-1. Human sera allowed for identification of endolysin epitopes frequently recognized by IgG in the human population. In some individuals, a relatively increased reactivity can be observed (Expanded View Figure EV4), although it is not clear whether this reactivity is due to natural contact with endolysins or it results from cross-reactions of antibodies induced by other unspecified antigens.

Antigenic epitopes were identified (Figure 3). To some extent, the epitopes identified with human sera were the same as those targeted by specific IgG developed in the animal model, particularly to epitopes in the C-terminal region of Cpl-1. However, other epitopes were not equally represented in both the human and mouse sera (Figure 2). This study included only natural antibodies in human populations, such as those induced by contact with epitopes that were the same or similar to the endolysin sequences or those antibodies that show cross-reactivity. However, the clinically relevant detection of immunoreactivity should include sera from patients treated with the investigated endolysins, hence our reliance on the mouse model sera rather than the human sera for identification of epitopes. This identification approach could be beneficial in future therapeutic trials, either for Pal and Cpl-1, or for other enzymes.

Identification of immunoreactive regions up to specific amino acids allowed for *in silico* design of new variants of the Pal and Cpl-1 endolysins. Our engineering approach was based on amino acid substitutions or deletion. Substitutions were chosen with the following assumptions: (i) amino acids that differed from the WT in their electrostatic charge and chemical structure were selected to disrupt epitope interaction with specific antibodies; (ii) the lowest energy state of the folded variant was preferred; (iii) smaller amino acids were preferred for the substitution due to lower steric tensions. Among the designed variants, only 3 out of 9 Pal variants were efficiently produced by the *E. coli* expression system in soluble forms. Five Cpl-1 variants, in contrast, were all successfully produced, although a lower yield was observed in the variant containing a triple-site substitution (Cpl-1 v4) and the deletion mutant (Cpl-1 v5). Not all variants retained good antibacterial activity (Figure 4), even though the selected amino acid substitutions were not within the catalytic site of the enzymes. These variants have limited, if any, value for the intended antibacterial applications. In the Cpl-1 variants, we observed that their poor antibacterial activity was linked to changes (either substitution or loss) of methionine at the 316^th^ position, as observed in variants v2, v4, and v5 (p>0.05 in comparison to WT Cpl-1 activity) (Figure 4, Table 1). This observation supports the key role of this amino acid in antibacterial activity of Cpl-1. Given that this mutation is in the cell wall binding domain, it likely affects binding to the pneumococcal surface.

Active enzymes can be prone to non-specific inactivation by serum components. If so, *in vivo* applicability of variants would be limited independent of their possible improved performance in the presence of specific antibodies. Thus, we first tested variants designed herein for their antibacterial activity *ex vivo* in naïve sera from unchallenged animals to identify potential non-specific inactivation. Notably, both WT enzymes were partially inactivated by non-specific serum and similar results were noted for the variants, with the activity of all enzymes in serum approximately 40% of their baseline activity in buffer. Nonetheless, three variants, Pal v3, Cpl-1 v1 and Cpl-1 v3, retained higher overall activity than the WT enzymes even in serum (Figure 5).

Since endolysin-specific antibodies from immunized animals decreased the rate of antibacterial activity of endolysins (Harhala et al., 2018; Rashel et al., 2007), we tested the new variants for their ability to escape immune responses by (i) lower efficacy of antibody induction, that is, by lower immunogenicity or (ii) by lower reactivity with antibodies induced by WT endolysins. Intrinsic immunogenicity of the variants was assessed in a mouse model and significant differences were only noted for two Pal variants, v3 and v9, although in the case of v3, its intrinsic immunogenicity was higher than that of WT Pal. Only in animals challenged with Pal v9 was the induced level of enzyme-specific IgG lower than in animals challenged with WT Pal. This effect was detectable only in the long-term observations on days 40 and 50 after challenge, but not at the earlier phase of response development (Figure 6). Thus, a partially deimmunized endolysin variant was successfully constructed by epitope engineering. The effect of deimmunization was, however, limited, since it was only observed in a later phase and the overall level of detected antibodies was significantly higher than that observed in non-challenged control mice. This suggests limited applicability of this particular variant as a deimmunized endolysin. Moreover, the Pal v9 variant demonstrated decreased intrinsic antibacterial activity due to the amino acid substitution, so its potential for therapeutic use can be expected to be weak. Nonetheless, our observation demonstrates that antigenic epitopes identified by epitope scanning can be deimmunized by substitutions of amino acids identified as key elements of the reactive epitope.

We further investigated cross-reactions of the Pal and Cpl-1 variants with IgGs induced by the WT enzymes, since escaping cross-reactions may thus be useful in repeat treatments. In all Pal variants, cross-reactivity was weaker than the reactivity of the WT Pal, and in Cpl-1, cross-reactivity was weaker in variants v4 and v5 (Figure 7). Since the overall bacteriolytic activity of Cpl-1 v4 and Cpl-1 v5 was low, we continued with Pal v3 and v9 in an *ex vivo* assay by blocking with WT Pal-specific serum. This assay models *in vivo* conditions in an organism where previous treatment with the WT endolysin yielded increased levels of specific IgG. No decrease of Pal v3 activity in WT Pal-specific serum was noted in comparison to its activity in naïve specific serum was observed. Thus, this variant escapes cross-reaction as well as cross-neutralization by specific antibodies induced by WT Pal. Furthermore, the overall antibacterial activity of Pal v3 in the presence of WT Pal-specific serum was significantly higher than of WT Pal in the same conditions (Figure 8). Two effects contributed to this advantage: Pal v3 was not neutralized by Pal-specific serum and its intrinsic activity was higher than Pal WT. Taken together, the data supports the use of Pal v3 in situations of repeated use by Pal where induction of antibodies may be a concern.

This study demonstrates a new efficient approach using EndoScan for identification of antigenic epitopes in proteins. EndoScan identifies physical interaction between protein-induced antibodies and the targeted protein might potentially complement epitope identification *in silico*. In recent years, a very interesting and promising approach for deimmunization of bacteriolytic proteins was demonstrated by Griswold and co-workers (Zhao et al., 2015, 2020), who used a computational method to design depletion of T cell epitopes in lysostaphin, an anti-staphylococcal enzyme. They demonstrated new variants of lysostaphin that could evade a specific immune response, but remain active against *Staphylococcus*. Here, we selectively modified epitopes identified with EndoScan in streptococcal endolysins and proposed new variants to decrease the immunoreactivity of these enzymes without losing antibacterial activity. To some extent, the immunogenicity of a protein can be decreased (as in Pal v9), or its cross-reactions with WT-induced antibodies can be minimized (as in Pal v3). Thus, EndoScan can be used for optimization of active enzymes designed for therapeutic applications, by modifying the immunogenicity of a protein without sacrificing its activity, with a relatively low number of tested proteins. Importantly, this approach is universal and can be applied to many active proteins, so its utility reaches far beyond endolysins and has potential in the design of numerous biological drugs.

## Conclusions

- Antigenic epitopes were identified in two endolysins, Pal and Cpl-1, by epitope scanning (EndoScan).
- Substitutions of key amino acids within the identified epitopes results in changes of protein immunogenicity, including identification of variants with lower immunogenicity.
- Substitutions of key amino acids within the identified epitopes may result in changes of protein cross-reactivity to specific IgG induced by the WT proteins.
- We propose Pal variant v3 (280-282: DKP→GGA) as a new engineered endolysin with higher antibacterial activity than WT Pal and capable of escaping cross-neutralization by antibodies induced by WT Pal.

## MATERIALS AND METHODS

### Epitope analysis

Identification of amino acids interacting with specific IgGs was performed in accordance with the protocol published by Xu et al. and adapted for our research using coding sequences of the investigated endolysins as the source for library design (Xu et al., 2015). The peptide library was created by selecting 56 amino acid (aa) long fragments tailing through the Cpl-1 (acc. no.: CAA87744.1) and Pal (acc. no: YP_004306947.1) endolysins, starting every 10 aa from the first amino acid, or where necessary, shorter oligopeptides near the end of the protein. In addition to the WT peptides, variants with consecutive single alanine substitutions, double alanine substitutions (i.e., targeted amino acid plus a neighboring amino acid), and triple alanine substitutions (i.e., targeted amino acid plus both neighboring amino acids) were also created. Alanine residues in the original sequences were substituted by glycine. Thus, our library contains all possible single-, double- and triple-substitutions for each position in the sequence for all oligopeptides designed as overlapping parts of the investigated endolysins.

The peptide library was reverse-translated into DNA sequences using codons optimized for expression in *E. coli*. The oligonucleotide library was synthesized using the SurePrint technology for nucleotide printing (Agilent). These oligonucleotides were used to create a phage library representing all oligopeptides using the T7Select 415-1 Cloning Kit (Merck Millipore). Immunoprecipitation of the library was performed in accordance with a previously published protocol (Harhala et al., 2022; Xu et al., 2015). Briefly, the phage library was amplified in a standard culture, purified by PEG precipitation, and dialyzed against Phage Extraction Buffer (20 mM Tris-HCl, pH 8.0, 100 mM NaCl, 6 mM MgSO_4_). All plastic equipment used for immunoprecipitation including Eppendorf tubes and 96-well plates were prepared by blocking with 3% bovine serum albumin (BSA) in TBST buffer overnight on a rotator (400 rpm, 4°C). A sample representing an average of 10^5^ copies of each clone was mixed with 2 μL of serum with high levels of anti-endolysin IgGs (see: specific sera induction in mice) or human serum from healthy volunteers. This mix was prepared in 250 μL phage extraction buffer and incubated overnight at 4°C on a rotator (400 rpm). A 40 μL aliquot of a 1:1 mixture of Protein A and Protein G Dynabeads (Invitrogen) was added and incubated for an additional 4 h at 4°C on a rotator (400 rpm). Liquid in all wells was separated from Dynabeads on a magnetic stand and removed. Beads were washed 3 times with 150 μL of a wash buffer (50 mM Tris-HCl, pH 7.5, 150 mM NaCl, 0.1% Tween-20) and beads were resuspended in 60 μL of water.

The immunoprecipitated part of the library was then used for amplification of the insert region according to the manufacturer’s instructions with a Phusion Blood Direct PCR Kit (Thermo Fisher Scientific). The primers T7_Endo_Lib_LONG_FOR GCCCTCTGTGTGAATTCT and T7_Endo_Lib_LONG_REV GTCACCGACACAAGCTTA were used and a second round of PCR was carried out with the IDT for Illumina UD indexes (Illumina Corp.) to add adapter tags. Sequencing of the amplicons in accordance with Illumina next generation sequencing (NGS) technology was outsourced (Genomed, Warszawa).

### Sequencing data analysis

Sequenced amplicons were mapped to the original nucleotide library sequences by the bowtie2 software as published by Xu et al. (Langmead & Salzberg, 2012; Xu et al., 2015). NGS sequencing reads were mapped by the bowtie2 software package with use of the file with list of oligonucleotide sequences synthesised originally to create a library as indexes (local mode)(Langmead et al., 2009; Langmead & Salzberg, 2012). The number of hits that mapped to each reference sequence (highest score) was counted (*count, c*).

The *signal* in each sample was calculated according to a formula (1).

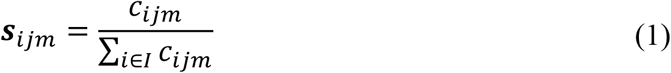

s – signal of i-th sequence in j-th serum sample and m-th technical replicate

c – number of reads mapped to i-th sequence in m-th technical replicate of j-th serum sample

I – set of all reference sequences (used as indexes to in mapping by the bowtie2 software)

The vast majority of clones were washed away and were not enriched, if detected at all. Thus, a zero-inflated negative binomial model was used to evaluate the probability of each signal to be random. Noteworthy, this model fitted control data better than zero-inflated Poisson distribution proposed by Xu et al. (Xu et al., 2015). Next, all p-values of oligopeptides detected in the sample were adjusted for multiple hypotheses (false discovery rate method) (Benjamini & Hochberg, 1995). *Input sample*: sample of phage library before immunoprecipitation representing starting levels of oligopeptides in the immunoprecipitation procedure. *Relative signal* (enrichment): an average signal in technical replicates of the sample divided by the average signal in ‘input samples’ of the same sequence is a signal ratio (*relative signal*). A series of t-tests evaluated the significance of differences in signals for each oligopeptide separately between input samples (one group) and the tested sera.

### Variant design

Every detected immunogenic amino acid was substituted *in silico* with every other possible amino acid and the differences in free folding energy (ΔΔG) was calculated using the FoldX software (Schymkowitz et al., 2005). The three-dimensional structure of Cpl-1 was downloaded from the PDB (Code: 2IXU) (Pérez-Dorado et al., 2007) and a model of the Pal structure was created using the I-Tasser software since its crystal structure has not previously been determined(Ambrish Roy et al., 2005). Structural variants were then chosen based on the following assumptions:

a. amino acids that differed from WT in their electrostatic charge and chemical structure were selected to disrupt epitope interaction with specific antibodies;
b. the lowest energy state of the folded variant was preferred;
c. smaller amino acids were preferred for the substitution due to lower steric tensions.

### Protein expression

The endolysins Pal (acc. no. YP_004306947) (Pal WT), derived from *Streptococcus* phage Dp-1, and Cpl-1 (acc. no. CAA87744) (Cpl-1 WT), derived from *Streptococcus* phage Cp-1, were used in this study. The Cpl-1 and Pal coding sequences were cloned into pBAD24 with a C-terminal 6xHis tag. Vectors with genes coding variants of Pal WT were synthesized *de novo* and cloned into the pBAD_HisA plasmid (BioCat GmbH, Germany).

Cpl-1 WT and Cpl variant 4 (v4) (see Table 1) were synthesized *de novo* and additional mutations were introduced by PCR with primers coding for the mutagenized site and ligated with Liga5 (A&A Biotechnology). Vector inserts were sequenced by Sanger sequencing to confirm validity of the cloning procedure. These vectors were transformed into *E. coli* B834(DE3) cells (EMD), and grown at 37°C with shaking in Luria-Bertani (LB) broth (10 g/L tryptone, 10 g/L NaCl, 5 g/L yeast extract) supplemented with ampicillin (50 mg/L) (Sigma-Aldrich, Europe), until the OD_600_ reached 1.0. Then, protein expression was induced by the addition of arabinose at a final concentration of 2.5 g/L (0.25%). The culture was incubated overnight at 22°C with intensive shaking.

### Protein purification

Bacteria were harvested using centrifugation (7 000 x g, 5 min) and suspended in PBS (140 mM NaCl, 2.68 mM KCl, 1.47 mM KH_2_PO_4_, 6.46 mM Na_2_HPO_4_, pH 7.2), which was supplemented with PMSF (1 mM) and lysozyme (0.5 mg/mL). The slurry was incubated for 6–7 h on ice and lysed using the freeze-thaw method. Mg^2+^ (up to 0.25 mM), DNAse (up to 20 μg/mL), and RNAse (up to 40 μg/mL) were then added to the extract and incubated on ice for 3 h. The fractions were separated using centrifugation (12 000 x g, 30 min, 4 °C) and the soluble fraction (supernatant) was collected.

For proteins to be used for mouse immunization studies, the crude supernatant was then incubated with NiNTA agarose (Qiagen) at room temperature, washed with PBS (5× volume of the agarose), and subsequently washed with an increasing concentration of imidazole (20 mM, 100 mM, 250 mM, and 500 mM). The 100 mM and 250 mM fractions containing the eluted endolysins were dialyzed against PBS at 4 °C and they were further purified using gel filtration (fast protein liquid chromatography) on a Superdex 75 10/300 GL column (GE Healthcare Life Sciences). The final step was LPS removal, which was performed with an EndoTrap Blue column (Hyglos GmbH, Munich, Germany). Purified, endotoxin-free protein samples were dialyzed against PBS and filtered through sterile 0.22-μm polyvinylidene difluoride filters (Millipore).

For proteins used in all other experiments, Cpl-1 and Pal (both WT and variants) were supplemented in PBS up to 75 mM before incubation with NiNTA agarose. Then the slurry was washed with 10x of 85 mM imidazole in PBS and the active protein was eluted with 250 mM imidazole and dialyzed three times against 100-fold excess of PBS at 4 °C (3,000 kDa molecular weight cut-off, Spectra/Por, Repligen). Finally, samples were filtered through sterile 0.22-μm polyvinylidene difluoride filters (Millipore). Samples were monitored by SDS-PAGE at all stages. Protein concentration was determined by the Bradford assay (Sigma-Aldrich, Europe) following the manufacturer’s instructions.

### Testing activity, fluorometric assay

The Sytox Green solution from the ViaGram Red+ Bacterial Gram Stain and Viability Kit (Thermo Fisher Scientific) was used to measure bacteriolytic activity via a fluorescence plate reader in accordance to a previously published protocol (Harhala et al., 2021). In short, mixtures of bacteria, endolysin, and Sytox™ Green (ThermoFisher Scientific) were prepared. Immediately after mixing, changes in the fluorescence signal of Sytox™ Green were measured for 15 min. The raw data from the fluorometric assay was normalized to the progress of the lysis with values between 0 (no lysis) and 1.0 (complete lysis). Progress values were used to calculate lytic activity as part of the total bacterial sample lysed per unit of time [min^-1^] using a linear regression model for Cpl-1 and a one-phase association model for Pal as previously described. Lytic activity is defined as the highest rate of lysis per unit of time for the linear part of the curve (for Cpl-1) or the beginning of the reaction (for Pal) before the reaction slows down due to diminishing bacterial substrate in the sample as described (Harhala et al., 2021).

### Specific sera induction in mice

C57BL/6J normal female mice (N=6 or 7) were obtained from Mossakowski Medical Research Centre Polish Academy of Sciences in Warsaw and housed in the Animal Breeding Centre of the Hirszfeld Institute of Immunology and Experimental Therapy (HIIET) in an isolated area in specific pathogen free conditions. To produce endolysin-specific serum, C57BL6/J mice were challenged intraperitoneally (i.p.) with Pal, Cpl-1, or their variants (0.1 mg per mouse) on day 0. Murine blood was collected into clotting tubes under anesthesia from the tail vein on day 28. Serum was separated from the blood using double centrifugation (2 250 x g for 5 min and then 10 000 x g for 10 min). The sera were stored at −20°C.

For comparison the immunogenicity of Pal and its variants, blood was collected as above every 3–5 days until at least day 50. Similarly, to compare immunogenicity of Cpl-1 and its variants, mice were challenged twice, on day 0 and at least 30 days later, since overall immunogenicity of Cpl-1 is lower than that of Pal (Harhala et al., 2018).

### Measurement of specific IgG antibody levels

MaxiSorp flat-bottom 96-well plates (Nunc, Thermo Scientific) were coated in sterile conditions overnight at 4°C with endolysins or PBS as a control using 100 μL per well at a concentration of 10 μg/mL, except for Pal v1 that demonstrated lower signal unless used at concentration 80 μg/mL (Expanded View Figure EV1). Subsequently, wells were washed five times with PBS and blocked for 30 min with SuperBlock Blocking buffer (Life Technologies Europe BV, Bleiswijk, Netherlands) at 150 μL per well and at room temperature. The solution was removed and the plates were washed five times with 0.05% Tween-20 (AppliChem GmbH, Darmstadt, Germany) in PBS at 100 μL per well. One hundred microliters per well of diluted serum (1:350 in PBS) was applied to the wells coated with endolysins. Each sample was investigated in five repeats. The plates were incubated at 37°C for 2 h, after which serum was removed and the plates were washed five times with 0.05% Tween-20 in PBS at 100 μL per well. One hundred microliters per well of diluted detection antibody (peroxidase-conjugated goat anti-mouse IgG (Jackson ImmunoResearch Laboratories) was applied to the plates and incubated for 1 h at room temperature in the dark. The antibody solution was removed and the plates were washed with PBS with 0.05% Tween 20 five times at 100 μL per well. TMB (50 μL) was used as a substrate reagent for peroxidase according to the manufacturer’s instructions (R&D Systems) and incubated for 30 min in the dark. Twenty-five microliters of 2N H_2_SO_4_ was added to stop the reaction and the absorbance was measured at 450 nm (main reading) and normalized by subtracting the background absorbance at 550 nm.

### Protein concentration measurement

Protein concentration was determined with use of the Bradford assay (Sigma) according to manufacturer’s manual if not explicitly mentioned otherwise. In short: 200μl of Bradford reagent was put in a well of a 96-well plate and 6.66 μl of the protein sample was added. After three min, the sample was mixed by pipetting and the absorbance was measured at 590nm and 450nm. The ratio was compared to a calibration curve (BSA in concentrations from 62.5 μg/mL to 1000 μg/mL) and the concentration of the protein in the sample was calculated from a linear regression of the standards. If the concentration of the protein was too high, the samples were diluted 4-fold.

### Ethical statement

All human subjects gave informed consent for inclusion before they participated in the study. The study was conducted in accordance with the Declaration of Helsinki, and the protocol was approved by the Bioethics Committee of the Regional Specialist Hospital in Wrocław (Project identification code: KB/nr 2/2017). All animal experiments were performed according to EU directive 2010/63/EU for animal experiments and were approved by the 1^st^ Local Committee for Experiments with the Use of Laboratory Animals, Wrocław, Poland (project no. 76/2011). The authors followed the ARRIVE (Animal Research: Reporting of *in vivo* Experiments) guidelines.

## ACKNOWLEDGEMENTS

This work was supported by the National Science Centre in Poland, grant no. UMO-2015/18/M/NZ6/00412 (granted to KD).

## AUTHOR CONTRIBUTIONS

MH - executed majority of the experiments, including library design, cloning, expression, purification, immunoprecipitation and analysis of immunoreactive epitopes, endolysin expression and purification, ELISA testing, and Sytox assay for endolysin activity, he conducted statistical analysis and participated in concluding results; KG, IR, ZK, PM, and JM – helped in endolysin production and phage display library construction, ELISA assays, Sytox assay, and in animal experiments, WB - contributed with graphics, DN - supervised all aspects of endolysin biology and biotechnology, participated in study design and in writing the manuscript, BO –helped in purification of samples for animal experimentation, KD-conceived and designed the study, executed and supervised animal studies and immunological aspects, analysed and concluded results, drafted the manuscript.

## Expanded View

**Expanded View Table EV1.**
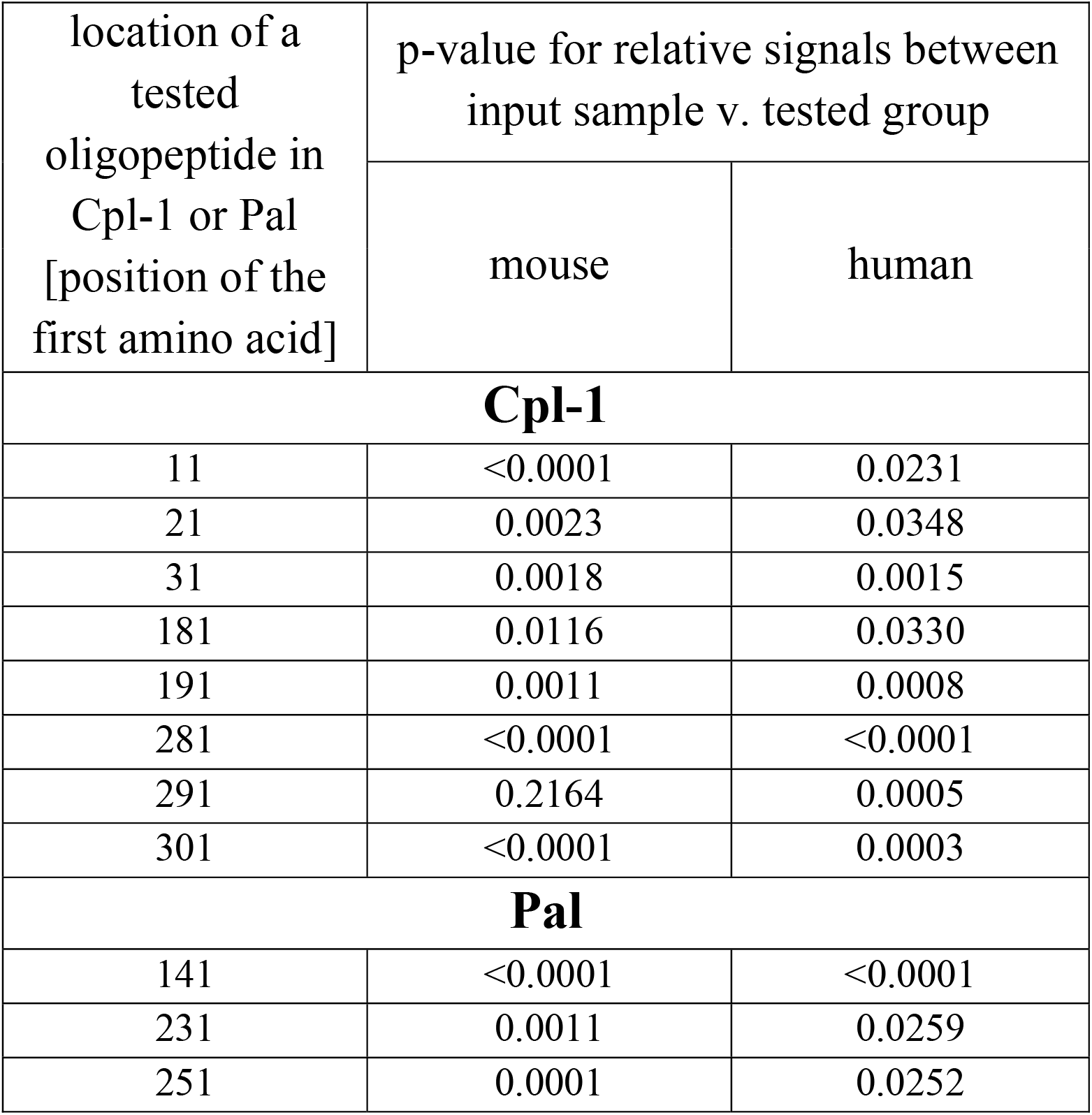
P-values for relative signals between the input sample v. tested group. Only adj. p-values < 0.05 are presented. Adj. p-values between relative signal (ratio) of a measured oligopeptide after immunoprecipitation and before (control level in input sample), values were calculated by GraphPad Prism 9 using the Kruskal Walis test, adjusted for multiple comparisons (Bemjamini, Krieger and Yekutieli). The control group is the library before immunoprecipitation.

| location of a tested oligopeptide in Cpl-1 or Pal [position of the first amino acid] | p-value for relative signals between input sample v. tested group |  |
| --- | --- | --- |
|  | mouse | human |
| **Cpl-1** |  |  |
| 11 | <0.0001 | 0.0231 |
| 21 | 0.0023 | 0.0348 |
| 31 | 0.0018 | 0.0015 |
| 181 | 0.0116 | 0.0330 |
| 191 | 0.0011 | 0.0008 |
| 281 | <0.0001 | <0.0001 |
| 291 | 0.2164 | 0.0005 |
| 301 | <0.0001 | 0.0003 |
| **Pal** |  |  |
| 141 | <0.0001 | <0.0001 |
| 231 | 0.0011 | 0.0259 |
| 251 | 0.0001 | 0.0252 |

**Expanded View Table EV2.** Statistical analysis of Pal and Cpl-1 variants’ cross-reactivity. Data information: Unpaired *t*-test between reactivity of specific IgG in shown groups, p-value adjusted using the Benjamini & Hochberg method, protocol proposed by Benjamini & Hochberg (1995) (Figure 2), 6 biological replicates in each group.

|  |  |
| --- | --- |
| Cpl-1 WT v. Cpl-1 v4 | p=0.0010 |
| Cpl-1 WT v. Cpl-1 v5 | p=0.00036 |
| Cpl-1 WT v. Cpl-1 v1 | p=0.016 |
| Cpl-1 WT v. Cpl-1 v2 | p=0.016 |
| Cpl-1 WT v. Cpl-1 v3 | p=0.0051 |
| Pal WT v. Pal v1 | p=0.0000024 |
| Pal WT v. Pal v3 | p=0.00000011 |
| Pal WT v. Pal v9 | p=0.00000018 |

**Expanded View Figure EV1.**
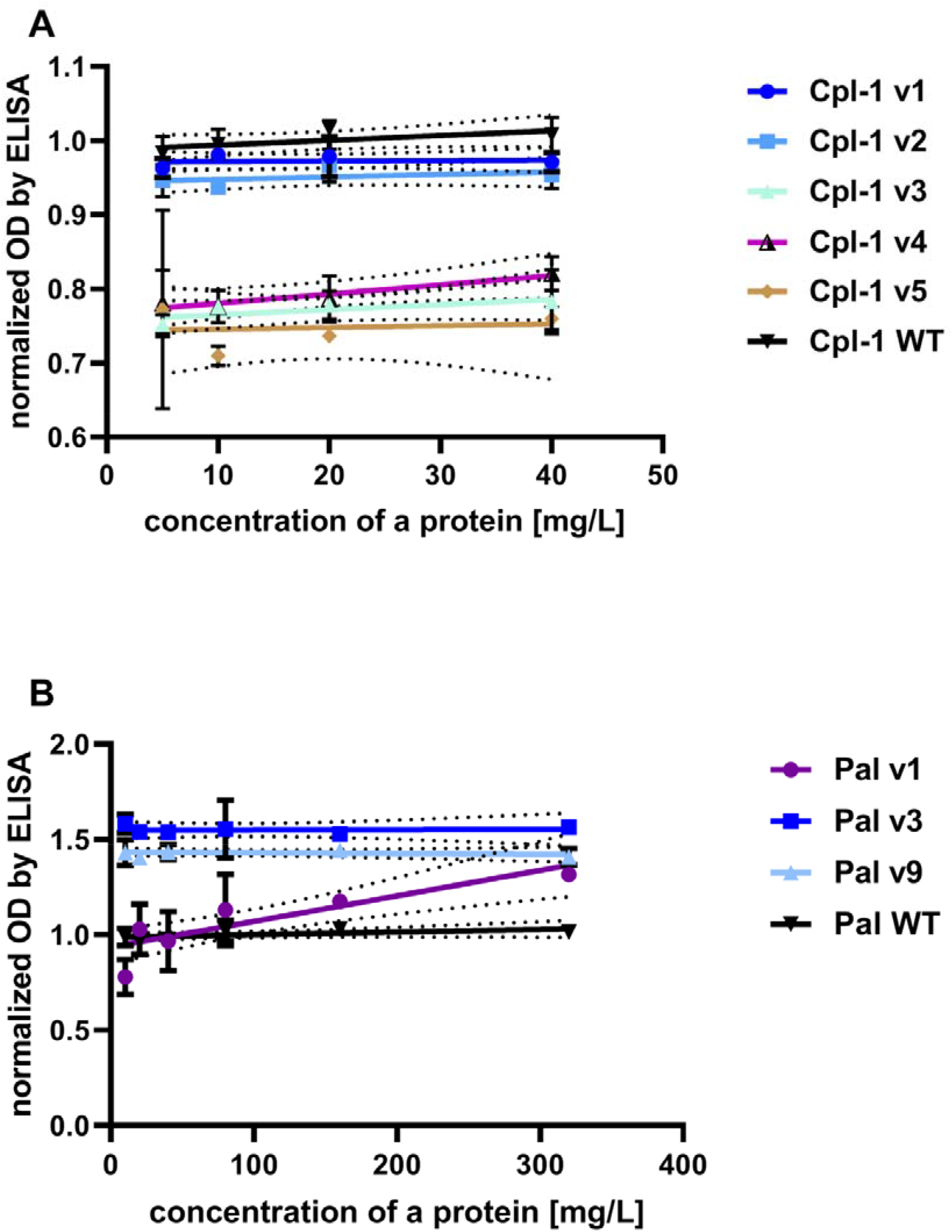
Levels of specific IgG in detected in wells coated with varying concentration of proteins. A, Normalized levels of specific IgG detected in wells coated with differing concentrations of Cpl-1 variants. B, Normalized levels of specific IgG detected in wells coated with differing concentrations of Pal variants. Data information: OD readings by ELISA are normalized to the average signal from WT protein set to 1.0. Points and whiskers represent mean and standard deviation of 6 biological replicates. Line represent linear regression model (dotted line represent 95% confidence interval) calculated and plotted by GraphPad Prism 9.

**Expanded View Figure EV2.**
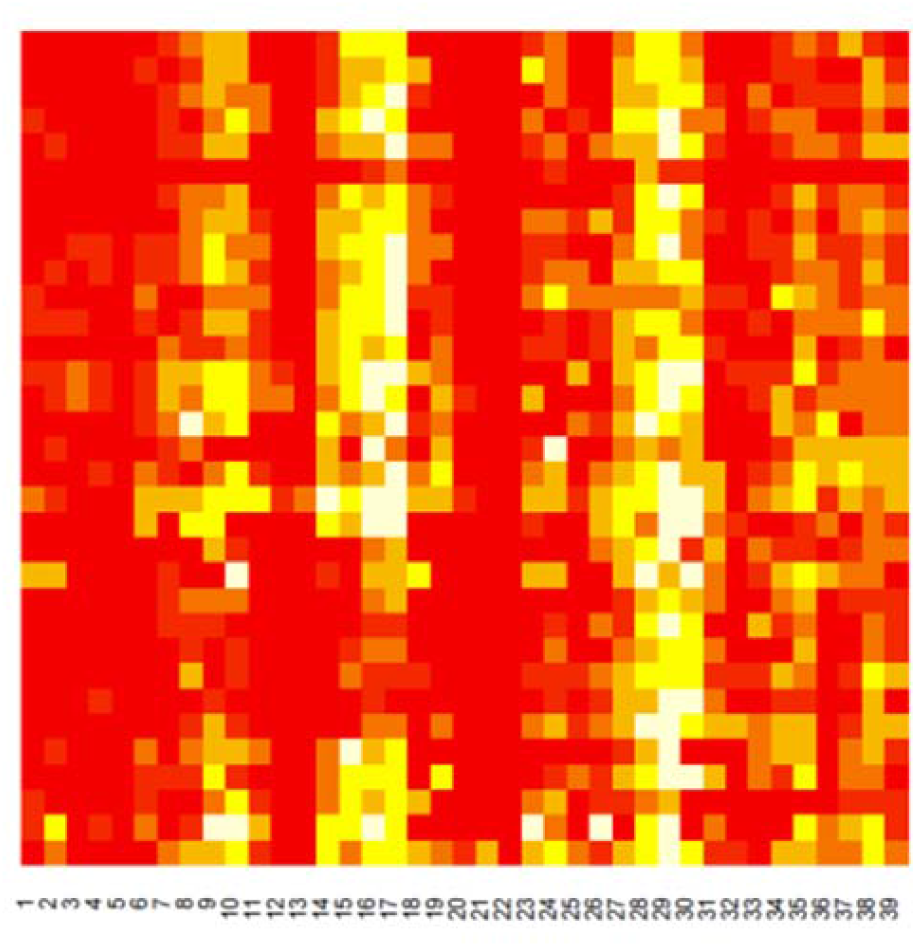
An examplary heatmap – graphical representation of data used for defining key amino acids within immunogenic regions (as demonstrated in Figure 3). Here for Cpl-1 – oligopeptide no. 31 starting after the 300^th^ amino acid position in the protein. Each row represent murine serum sample, each column represent oligopeptide no 31 with a specific (number on x-axis) amino acid substituted with alanine (or glycine if it previously was an alanine). Colors show the ratio of detected signal between alanine-substituted and unsubstituted oligopeptides (mean of 6 biological replicates). White square means a ratio below 20%, red square show a ratio over 90% (no signal lost by substitution).

**Expanded View Figure EV3.**
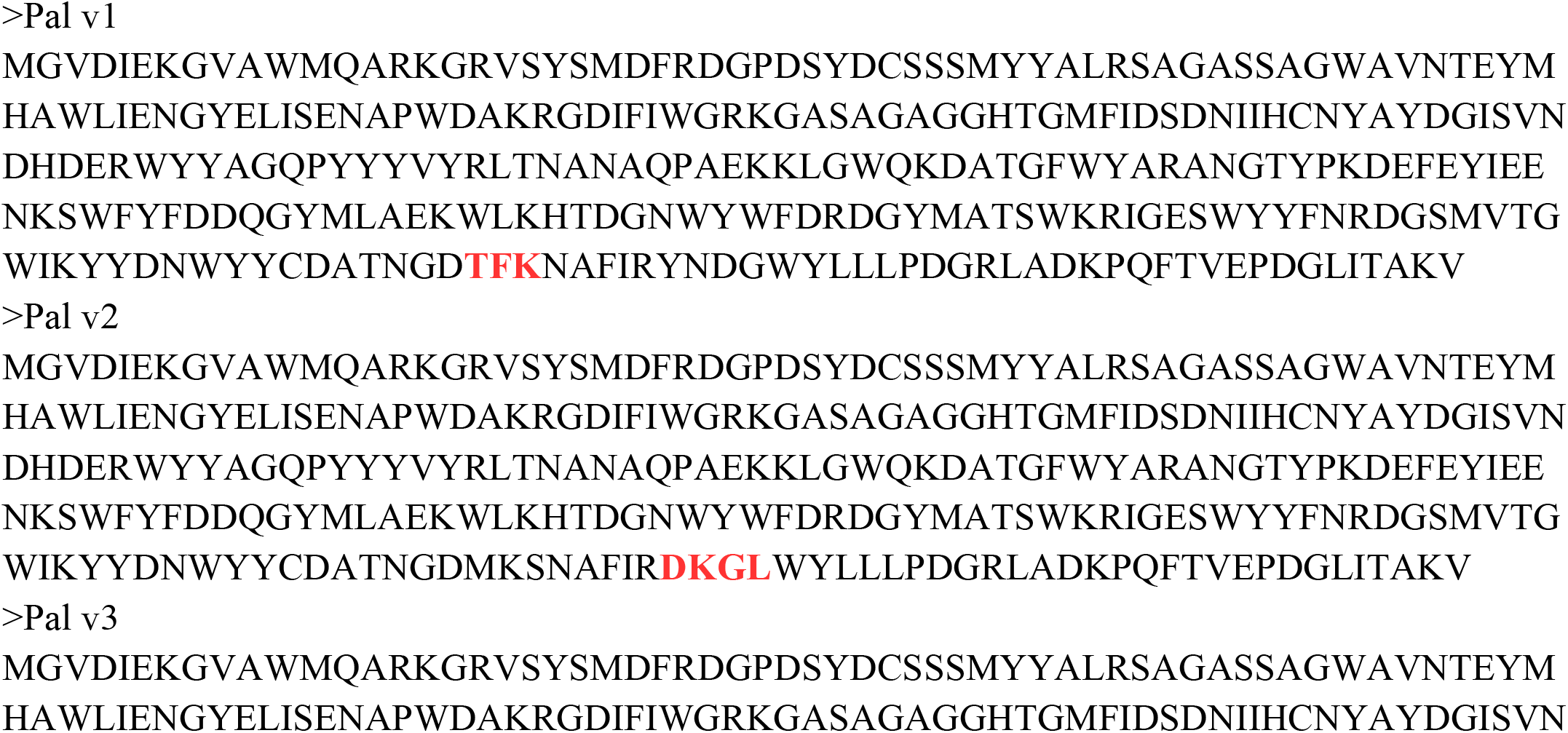
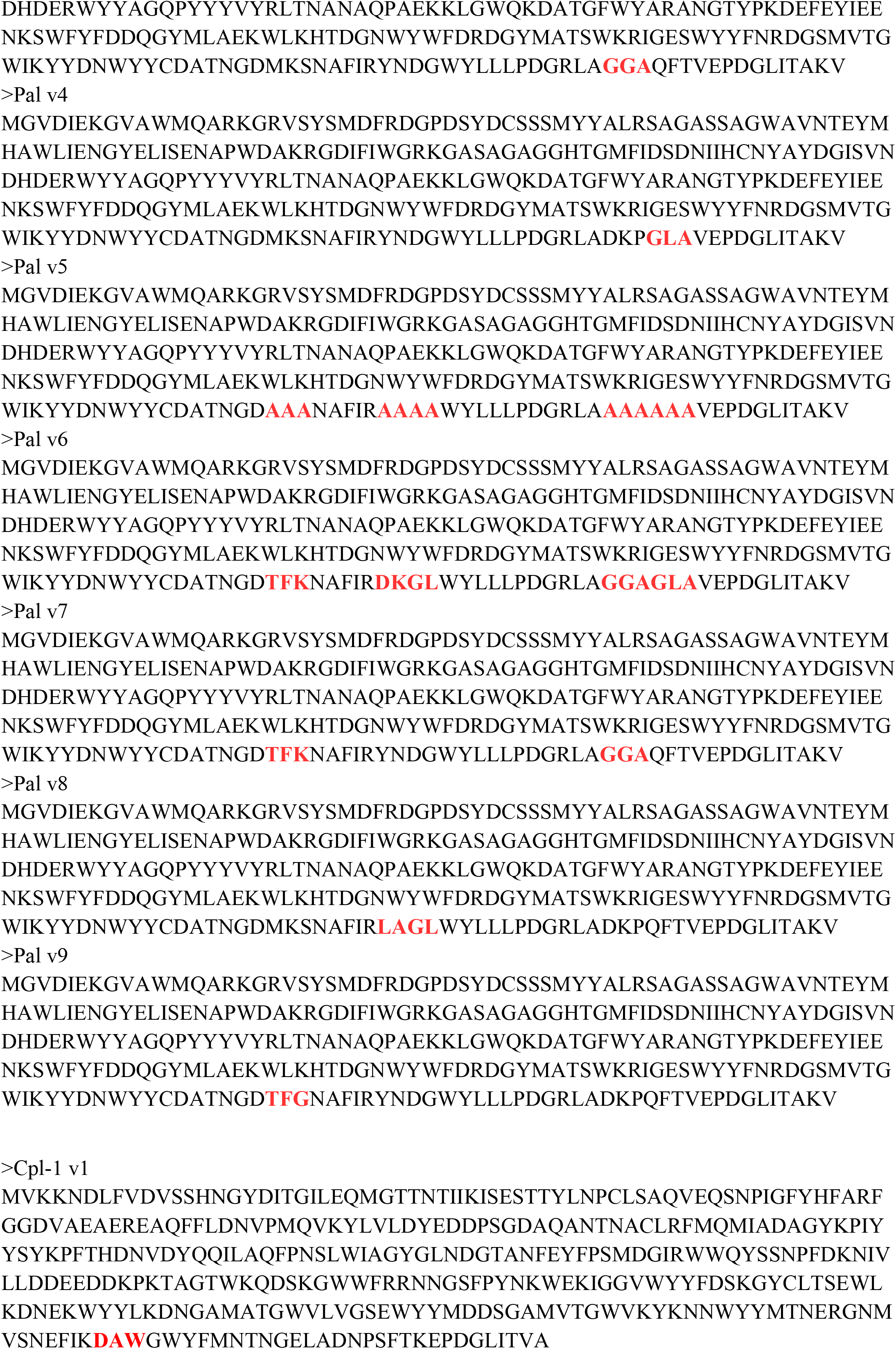
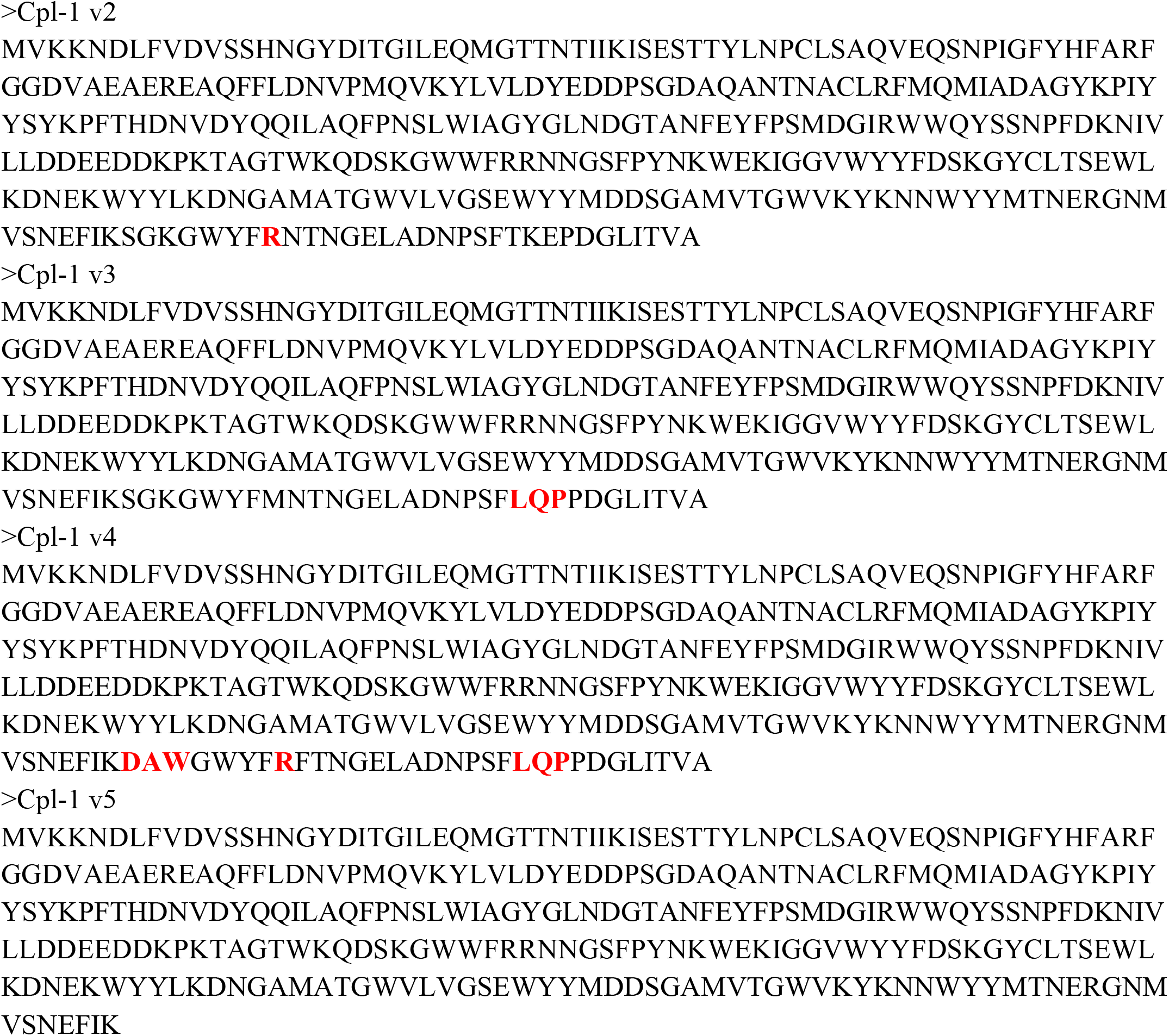
Full list of variants with modified epitopes identified by EndoScan in fasta format.

**Expanded View Figure EV4.**
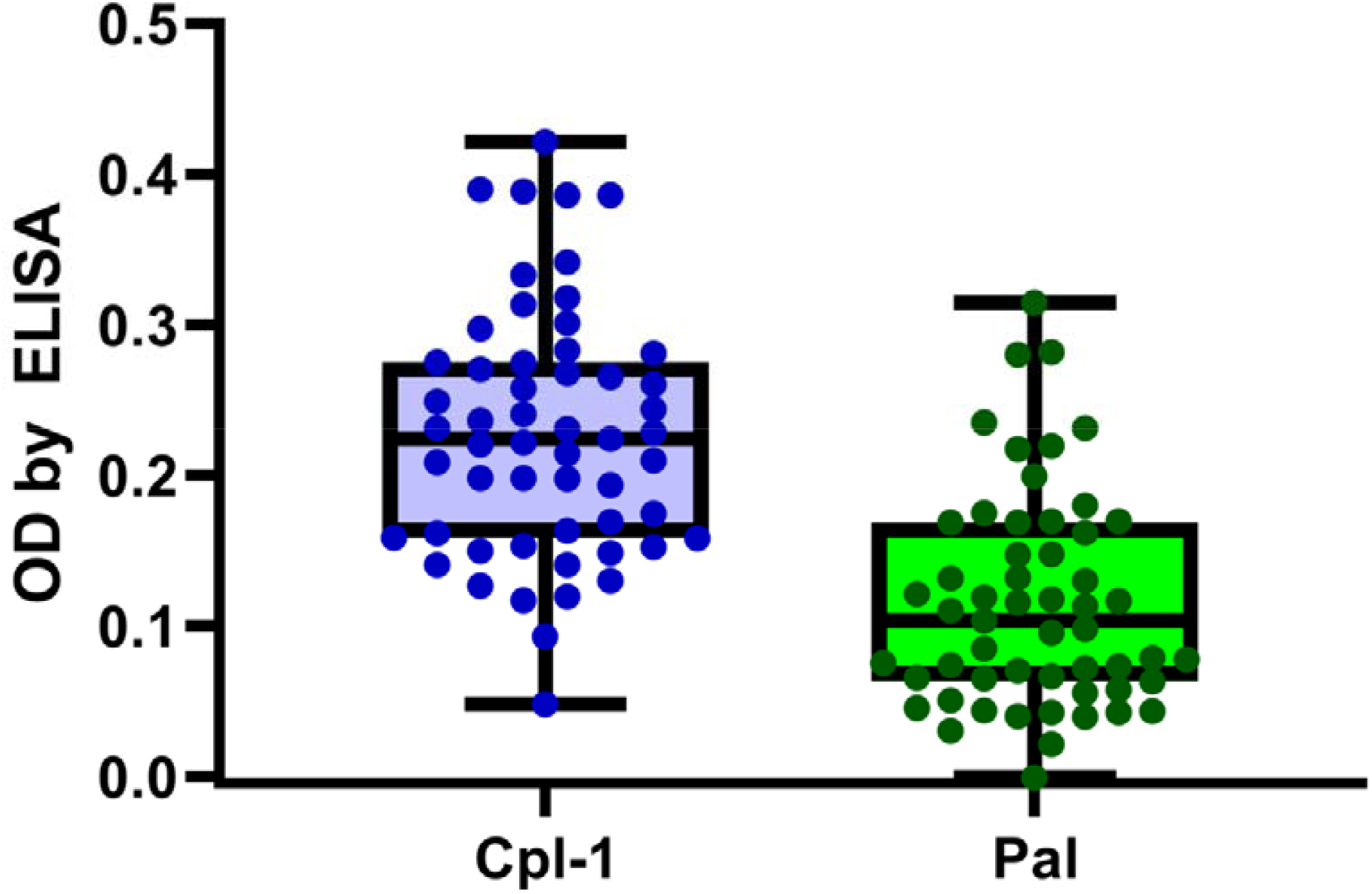
Distribution of endolysin-specific antibodies in humans. Data information: European population (N=56); dots represent individual samples values (mean of 3 technical replicates); line and whiskers represent median and min or max read respectively. Upper and lower border of box represent 75^th^ and 25^th^ quantile of data. One serum sample showed an abnormally high reading against both proteins (>0.6), resulting in this sample being identified as an outlier (ROUT method, GraphPad Prism 9, cut-off - 1%) and it’s not plotted.

